# Deconvoluting Latent Erythroid–Mitochondrial Transcriptomic Gradients and Technical Sample Structure in Severe Preeclampsia: A Stratified Sensitivity Framework for Public Placental RNA-Seq Datasets

**DOI:** 10.64898/2026.11.20.768086

**Authors:** Khondoker Ashab Zaman, Taufiq Mustafizur Rahman, Md Shariful Hasan, Khondoker Qumruzzaman, Umme Kulsum

## Abstract

**Background:** Bulk placental transcriptomic profiling has been widely deployed to elucidate the molecular pathogenesis of severe preeclampsia (sPE) and to nominate clinical biomarkers across diverse maternal populations. However, human placental tissue exhibits profound cellular heterogeneity and vascular perfusion variability. When unmeasured technical processing structure or erythroid–trophoblast compositional gradients covary with clinical phenotypes in public repositories, standard differential gene expression (DGE) pipelines risk severe false-discovery inflation and spurious ancestry-specific biomarker claims.

**Methods:** We developed and applied a multi-layered transcriptomic audit, variance-partitioning diagnostic, and stratified permutation framework to re-evaluate the public NCBI Gene Expression Omnibus dataset GSE234729 (14,670 filtered, DESeq2-normalized features across 123 columns). Following strict provenance verification that excluded 12 unverified cross-series columns (linked to GSE186257), we analyzed 111 verified placental transcriptomes (43 sPE and 68 normotensive controls across African, Asian, and European maternal ancestries). Expression values were modeled on the log2(E+1) scale using robust empirical Bayes linear modeling (limma; ~ group + ancestry + fetal sex), benchmarked against standard empirical Bayes implementations, principal component variance decompositions, erythroid and mitochondrial gene-set scoring, stratified fixed-effect inverse-variance meta-analysis with Cochran’s Q heterogeneity testing, and 1,000-iteration structure-preserving Monte Carlo permutation schemes.

**Results:** While standard primary modeling identified 5,667 differentially expressed features at Benjamini–Hochberg false discovery rate (FDR) < 0.05 (5,680 under standard empirical Bayes), unsupervised principal component analysis revealed that the dominant axis of transcriptomic variation (PC1, explaining 38.2% of total variance; 31.9%–47.3% across scaling/feature subsets) was orthogonal to disease biology. Instead, PC1 captured an extreme reciprocal gradient between haemoglobin genes (HBA1, HBA2, HBB, HBG1, HBG2; r = 0.97) and 16 mitochondrial (MT-) oxidative phosphorylation transcripts (r = −0.96). Critically, PC1 bifurcated the cohort in exact 100% correspondence with an undocumented sample-identifier prefix regime (58 samples labeled ‘P’ vs. 53 labeled ‘P1/P3/P6/P7/P8/P9’, accounting for 95.4% of PC1 variance). This latent prefix structure was severely entangled with clinical phenotype (57% sPE in non-‘P’ vs. 22% in ‘P’; Fisher’s exact P < 0.001) and maternal ancestry (15 of 16 African-ancestry samples confined to the non-‘P’ stratum). Adjusting for the ID-prefix regime reduced genome-wide significant calls by 35.9% (from 5,680 to 3,643 features), while structure-preserving permutations (Scheme B) demonstrated that covariate imbalance alone generated a median of 3,757 false-positive calls (95% interval: 1,423–5,617; empirical P = 0.023) in the complete absence of biological disease signal. Despite this pervasive genome-wide confounding, canonical syncytiotrophoblast anti-angiogenic and metabolic stress signatures—FLT1 (+2.45 log2 units, FDR = 3.9e-15), LEP (+4.91, FDR = 8.2e-13), PHYHIP (+1.42, FDR = 1.5e-10), ENG (+1.61, FDR = 3.4e-15), PAPPA2 (+2.02, FDR = 2.1e-11), FSTL3 (+2.65, FDR = 1.8e-08), and PPP1R1C (+1.56, FDR = 1.5e-10)—exhibited remarkable directional invariance across all sensitivity models and within-stratum meta-analyses (Cochran’s Q P > 0.39). Conversely, the putative cardiovascular risk biomarker IL3RA showed null primary association (−0.07 log2 units; 95% CI: −0.51 to 0.36; P = 0.74; genome-wide rank 13,299/14,670), reversed algebraic sign across strata (+0.77 in non-‘P’ vs. −0.54 in ‘P’; heterogeneity P = 0.003), and failed genome-wide ancestry interaction testing (FDR ≈ 1.0).

**Conclusions:** Nearly two-thirds of the apparent genome-wide differential expression in GSE234729 can be recapitulated by severe collinearity between clinical group, maternal ancestry, and a cryptic erythroid– mitochondrial sample structure. While core anti-angiogenic preeclampsia pathways (FLT1, LEP, ENG, PAPPA2, FSTL3, PHYHIP, PPP1R1C) are robust against this confounding, previously highlighted associations for IL3RA are statistically fragile and artifactually driven by unmodeled stratum heterogeneity. We establish an actionable bioinformatic auditing protocol requiring mandatory erythroid-contamination diagnostics, identifier-structure screening, and structure-preserving permutation testing prior to clinical biomarker nomination from public placental transcriptomes.

## 1. Introduction

Preeclampsia is a life-threatening multisystem hypertensive disorder of human pregnancy that complicates 2% to 8% of gestations globally and remains a leading driver of maternal and perinatal morbidity and mortality, particularly in low- and middle-income settings and among historically underserved racial and ethnic populations [1,2]. Pathophysiologically, severe preeclampsia (sPE) is anchored in defective early extravillous trophoblast invasion and incomplete spiral artery remodeling, which precipitate uteroplacental malperfusion, hypoxia–reoxygenation injury, syncytiotrophoblast endoplasmic reticulum and mitochondrial stress, and the systemic release of anti-angiogenic and pro-inflammatory factors into the maternal circulation [1,3]. Over the past decade, high-throughput bulk RNA sequencing (RNA-seq) of delivered placental villous tissue has served as the primary discovery engine for mapping the transcriptional architecture of sPE, identifying dysregulated modules across angiogenic signaling, trophoblast invasion, metabolic reprogramming, and maternal–fetal immune tolerance [2,4].

Recently, Aisagbonhi et al. [1] profiled bulk placental transcriptomes from 123 deliveries (reported as 50 sPE cases and 73 normotensive controls) and deposited the processed expression data in the NCBI Gene Expression Omnibus (GEO) under accession GSE234729 [2]. Their study highlighted elevated placental expression of fms-related receptor tyrosine kinase 1 (FLT1), leptin (LEP), phytanoyl-CoA 2-hydroxylase interacting protein (PHYHIP), and interleukin-3 receptor subunit alpha (IL3RA) in individuals of African ancestry with sPE, proposing placental IL3RA upregulation as a novel molecular link to long-term maternal cardiovascular morbidity [1,2]. Because public transcriptomic datasets in NCBI GEO serve as foundational reference resources for secondary meta-analyses, machine-learning classifier training, and clinical biomarker prioritization, the statistical validity of their underlying expression signals is of paramount importance to the reproductive biology and translational bioinformatics communities.

However, bulk placental RNA-seq datasets are uniquely vulnerable to technical and biological confounding that can severely distort standard differential gene expression (DGE) pipelines [3,5]. First, the human placenta is a highly vascularized, cellularly heterogeneous organ comprising multinucleated syncytiotrophoblasts, villous cytotrophoblasts, extravillous trophoblasts, Hofbauer macrophages, fetal endothelial cells, and varying volumes of retained maternal and fetal blood [6,7]. Variations in tissue harvesting depth, PBS washing stringency, exsanguination protocols, or mode of delivery can introduce dramatic sample-to-sample fluctuations in retained erythrocytes and reticulocytes, which carry massive quantities of globin mRNAs (e.g., HBA1, HBA2, HBB, HBG1, HBG2) alongside depleted nuclear and mitochondrial transcripts [6,7]. Second, when multi-ethnic cohorts are assembled across sequential clinical recruitment waves, hospital sites, or RNA extraction and sequencing batches without explicit batch tracking in public metadata, latent technical axes frequently become entangled with clinical group and maternal ancestry [3,8]. When case–control ratios or ancestral subgroups are unevenly distributed across a dominant non-biological axis of variance, conventional linear models yield thousands of spurious false-positive genes that reflect sample collection architecture rather than disease pathophysiology [3,8,9].

Despite the widespread reuse of GEO expression matrices, systematic forensic auditing of sample-level provenance, identifier-linked latent structure, and structure-preserving null distributions is rarely performed before secondary biomarker claims are propagated. In the present study, we execute a comprehensive methodological re-evaluation of GEO dataset GSE234729. Rather than treating the deposited matrix as a uniform biological cohort, we establish a multi-tiered diagnostic and sensitivity framework designed to: (i) audit sample-to-metadata provenance and resolve cross-series contamination; (ii) decompose the dominant axes of transcriptomic variance into biological versus erythroid– mitochondrial and identifier-linked structural components; (iii) quantify the exact inflation of genome-wide false discoveries attributable to group-by-structure imbalance using novel stratified Monte Carlo permutation schemes; and (iv) rigorously adjudicate the stability of canonical sPE genes (FLT1, LEP, PHYHIP, ENG, PAPPA2, FSTL3, PPP1R1C) against the putative ancestry-associated biomarker IL3RA. Our findings provide both a definitive correction to the evidentiary interpretation of GSE234729 and a generalizable methodological blueprint for auditing public bulk transcriptomic repositories.

## 2. Materials and Methods

### 2.1. Data Provenance Audit, Matrix Curation, and Cohort Reconciliation

The processed placental gene expression matrix (GSE234729_Plac_samples_diff_ancestry_Deseq2_norm_filtered.txt.gz) and accompanying series matrix metadata were retrieved directly from NCBI GEO under accession GSE234729 [2]. Inspection of the deposited file revealed 14,670 gene features across 123 sample columns containing non-negative, non-integer values pre-normalized via DESeq2 size-factor scaling. Because raw integer read counts and FASTQ files were not deposited in the public repository, count-based negative binomial generalized linear modeling (such as native DESeq2 or edgeR quasi-likelihood pipelines) could not be validly applied.

A rigorous sample-provenance audit was conducted by cross-referencing the 123 matrix column headers against the official GSM sample records registered under series GSE234729. Exactly 111 matrix columns matched verified GSE234729 sample records. The remaining 12 columns lacked corresponding GSM entries within GSE234729; forensic cross-series tracking in our reproducible workflow linked these 12 identifiers and their near-identical expression profiles to an earlier, related GEO submission (GSE186257). Because independent clinical verification of maternal ancestry, diagnostic criteria, and covariates was unavailable for these 12 unlinked columns within the GSE234729 registry, they were strictly excluded from all primary and sensitivity models to prevent cross-series verification bias. The final reconciled analytic cohort comprised N = 111 verified placental samples (43 severe preeclampsia cases and 68 normotensive non-hypertensive controls; Table 1). Furthermore, structural auditing of feature annotations revealed 733 duplicated HUGO gene symbols within the 14,670-row matrix; consequently, all computational indexing, modeling, and multiple-testing corrections were executed using unique row positions and Ensembl gene identifiers (ENSG) prior to symbol mapping.

**Table 1.** Demographic and clinical distribution of the provenance-verified GSE234729 placental transcriptomic cohort (N = 111).

| Clinical Group | Total (n) | African Ancestry | Asian Ancestry | European Ancestry | Female Fetus | Male Fetus |
|---|---|---|---|---|---|---|
| Severe Preeclampsia (sPE) | 43 | 6 (14.0%) | 17 (39.5%) | 20 (46.5%) | 19 (44.2%) | 24 (55.8%) |
| Normotensive Control | 68 | 10 (14.7%) | 14 (20.6%) | 44 (64.7%) | 40 (58.8%) | 28 (41.2%) |
| **Total Verified Cohort** | **111** | **16 (14.4%)** | **31 (27.9%)** | **64 (57.7%)** | **59 (53.2%)** | **52 (46.8%)** |

*Note: Values represent sample counts (n) and row percentages. Twelve additional matrix columns lacking GSE234729 sample records (traced to GSE186257) were excluded prior to modeling.*

### 2.2. Clinical Covariates and Unmeasured Confounder Assessment

Phenotypic variables extracted from the verified GEO sample records comprised clinical diagnostic group (severe preeclampsia vs. normotensive control), self-reported maternal racial/ethnic ancestry (categorized as African, Asian, or European), and fetal sex (female vs. male; the biological or clinical basis of fetal sex assignment was not specified in the GEO metadata). Critical obstetric and technical covariates known to influence placental gene expression—specifically gestational age at delivery, maternal pre-pregnancy body mass index (BMI), neonatal birth weight, mode of delivery (cesarean section vs. vaginal delivery), presence of labor, placental anatomical sampling site, tissue washing protocol, RNA integrity number (RIN), and library preparation or sequencing run batch—were absent from the public metadata record.

### 2.3. Primary Differential Expression Modeling and Empirical Bayes Estimation

To stabilize variance across the dynamic range of non-integer DESeq2-normalized expression values (E), the matrix was transformed onto the log2(E + 1) scale. The prespecified primary linear model was parameterized as:

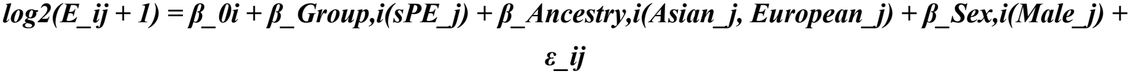

yielding a full-rank 5-column design matrix with 106 residual degrees of freedom across the 111 samples. Linear modeling was executed in R using the limma framework (lmFit) followed by empirical Bayes variance moderation (eBayes) with robust hyperparameter estimation enabled (robust = TRUE) to protect against hypervariable outlier genes and preserve statistical power [4–6]. Model coefficients (β_Group) represent covariate-adjusted differences in log2(E + 1) units between severe preeclampsia and normotensive controls rather than simple ratio fold changes. Genome-wide multiplicity was controlled using the Benjamini–Hochberg (BH) step-up procedure [7] across all 14,670 features, with statistical significance defined at an exploratory threshold of FDR < 0.05. Feature-level standard errors (SE) were recovered exactly as the adjusted difference divided by the moderated t-statistic, and 95% confidence intervals (CIs) were computed using the model’s total empirical Bayes degrees of freedom (df_total = 108.7). To ensure complete methodological transparency, a standard (non-robust, robust = FALSE) empirical Bayes fit of the identical design matrix was also evaluated, yielding 5,680 significant features compared with 5,667 under the primary robust implementation; counts from the two variance-moderation implementations were strictly segregated in all downstream comparisons.

### 2.4. Unsupervised Variance Decomposition and Erythroid–Mitochondrial Diagnostics

To interrogate latent sample structure without phenotypic supervision, principal component analysis (PCA) was performed via singular value decomposition on mean-centered log2(E + 1) matrices across all 14,670 features, and systematically cross-checked using the top 500 and top 1,000 highest-variance features both with and without unit-variance scaling. To evaluate whether the dominant principal component (PC1) reflected biological tissue composition or technical processing gradients, sample-level composite expression scores were computed for two orthogonal physiological gene sets: (i) an erythroid/haemoglobin signature comprising the mean log2(E + 1) expression of five major adult and fetal globin genes (HBA1, HBA2, HBB, HBG1, and HBG2), indexing retained maternal and fetal blood content; and (ii) a mitochondrial oxidative phosphorylation signature comprising the mean log2(E + 1) expression of 16 mitochondrial genome-encoded (MT-) transcripts. Furthermore, we audited sample alphanumeric identifiers by parsing the prefix preceding the first underscore or numeric digit, which segregated samples into seven discrete string classes (P, P1, P3, P6, P7, P8, and P9) and a binary structural dichotomy (‘P’ prefix [n = 58] vs. non-‘P’ numbered prefixes P1/P3/P6/P7/P8/P9 [n = 53]). Because the original investigators provided no documentation linking sample ID prefixes to laboratory batches, we treat the binary ‘ID-prefix group’ strictly as an empirical descriptor of latent identifier-associated structure rather than a confirmed technical batch variable.

### 2.5. Multi-Model Sensitivity Architecture and Stratified Meta-Analysis

To quantify the sensitivity of both genome-wide discovery counts and candidate biomarker effect sizes to latent sample structure, five structured sensitivity models were executed using a unified standard moderated-t implementation (which exactly reproduces the baseline standard-eBayes count of 5,680 features): (Model S1) an extended linear model incorporating the binary ID-prefix group as an additive covariate (~ group + ancestry + fetal sex + ID_prefix_group); (Model S2) direct estimation of the ID-prefix group contrast itself, adjusted for ancestry and fetal sex; (Model S3) an eigen-gene adjusted model incorporating the first two principal components (~ group + ancestry + fetal sex + PC1 + PC2), recognizing that PC1 and PC2 are data-derived covariates partially collinear with disease status; (Model S4) a fully stratified analysis in which the primary linear model (~ group + ancestry + fetal sex) was fitted independently within the ‘P’ prefix stratum (n = 58) and the non-‘P’ prefix stratum (n = 53), followed by fixed-effect inverse-variance weighted meta-analysis and Cochran’s Q test for inter-stratum effect-size heterogeneity; and (Model S5) a joint 2-degree-of-freedom group-by-ancestry multiplicative interaction model to test whether candidate gene differences varied significantly across African, Asian, and European maternal ancestries.

### 2.6. Structure-Preserving Monte Carlo Permutation Framework

To determine how many apparently significant differentially expressed genes could arise purely from the unequal distribution of sPE cases across the latent ID-prefix structure under a true biological null hypothesis, we designed two 1,000-iteration Monte Carlo permutation experiments (fitting the baseline model ~ group + ancestry + fetal sex without an ID-prefix term at each iteration). In Permutation Scheme A (randomization seed = 20261030), clinical group labels (sPE vs. control) were randomly shuffled within joint ancestry × fetal sex strata, breaking the association between disease status and ID-prefix group while preserving demographic balance. In Permutation Scheme B (randomization seed = 20261031), clinical group labels were permuted strictly within joint ancestry × fetal sex × ID-prefix group three-way strata. Scheme B eliminates all true within-stratum biological differences between sPE and controls while exactly preserving the observed empirical confounding between sPE prevalence and the ID-prefix transcriptomic gradient, thereby yielding an exact empirical distribution of false-positive feature counts driven by structural imbalance alone.

### 2.7. Candidate Biomarker Adjudication and Reproducibility Standards

Eight candidate genes were subjected to granular effect-size, confidence-interval, and heterogeneity tracking across all modeling tiers. Four genes—FLT1, LEP, PHYHIP, and IL3RA—were prespecified directly from the title and primary claims of the original publication [1]. Four additional well-established preeclampsia genes—endoglin (ENG), pappalysin-2 (PAPPA2), follistatin-like 3 (FSTL3), and protein phosphatase 1 regulatory inhibitor subunit 1C (PPP1R1C)—were selected as data-informed positive reference controls after observing their prominent ranking in initial differential expression outputs (and confirming their secondary reporting in [1]). Minimum detectable effect sizes (MDES) at 80% statistical power (α = 0.05, two-sided) were calculated from the empirical Bayes standard errors. All analyses were executed in R (version 4.6.1) using limma (version 3.68.5) via an end-to-end automated script (GSE234729_FINAL_REPRODUCIBLE_ANALYSIS.R).

## 3. Results

### 3.1. Provenance Reconciliation and Demographic Architecture of the Verified Cohort

Cross-referencing the 123 columns of the deposited GSE234729 expression matrix against official NCBI GEO sample metadata confirmed 111 verified patients (43 sPE and 68 normotensive controls) and identified 12 unlinked columns originating from series GSE186257, which were excluded. As detailed in Table 1, the verified cohort was predominantly of European (n = 64; 57.7%) and Asian (n = 31; 27.9%) self-reported maternal ancestry, whereas the African-ancestry subgroup—the central focus of the original report’s title and biomarker claims—was severely limited in sample size, comprising only 16 total deliveries (14.4%), including merely 6 sPE cases and 10 controls. Fetal sex was balanced overall (59 female, 52 male fetuses), with a modest excess of male fetuses among sPE deliveries (55.8% vs. 41.2% in controls).

### 3.2. Baseline Genome-Wide Differential Expression and Candidate Biomarker Performance

When analyzed via the prespecified primary model (limma with robust empirical Bayes moderation adjusted for maternal ancestry and fetal sex), 5,667 of the 14,670 placental features (38.6% of the measured transcriptome) reached genome-wide significance at BH FDR < 0.05 (5,680 features under standard non-robust empirical Bayes). Examination of the eight focal candidate genes (Table 2) revealed highly significant upregulation in sPE across all seven canonical preeclampsia markers: FLT1 (+2.45 log2(E+1) units; t = 10.84; FDR = 3.9e-15), LEP (+4.91; t = 9.65; FDR = 8.2e-13), PHYHIP (+1.42; t = 8.42; FDR = 1.5e-10), ENG (+1.61; t = 11.00; FDR = 3.4e-15), PAPPA2 (+2.02; t = 8.92; FDR = 2.1e-11), FSTL3 (+2.65; t = 7.19; FDR = 1.8e-08), and PPP1R1C (+1.56; t = 8.43; FDR = 1.5e-10). In stark contrast, IL3RA exhibited complete absence of differential expression between sPE and normotensive controls in the primary adjusted model, yielding a near-zero negative point estimate of −0.07 log2(E+1) units (95% CI: −0.51 to 0.36; moderated t = −0.33; P = 0.740; BH FDR = 0.817), ranking 13,299th out of 14,670 genes. Based on its empirical Bayes standard error (SE = 0.22), our primary model possessed 80% statistical power (at two-sided α = 0.05) to detect a true group difference of approximately 0.62 log2(E+1) units, ruling out any moderate-to-large global elevation of IL3RA in sPE.

**Table 2.**
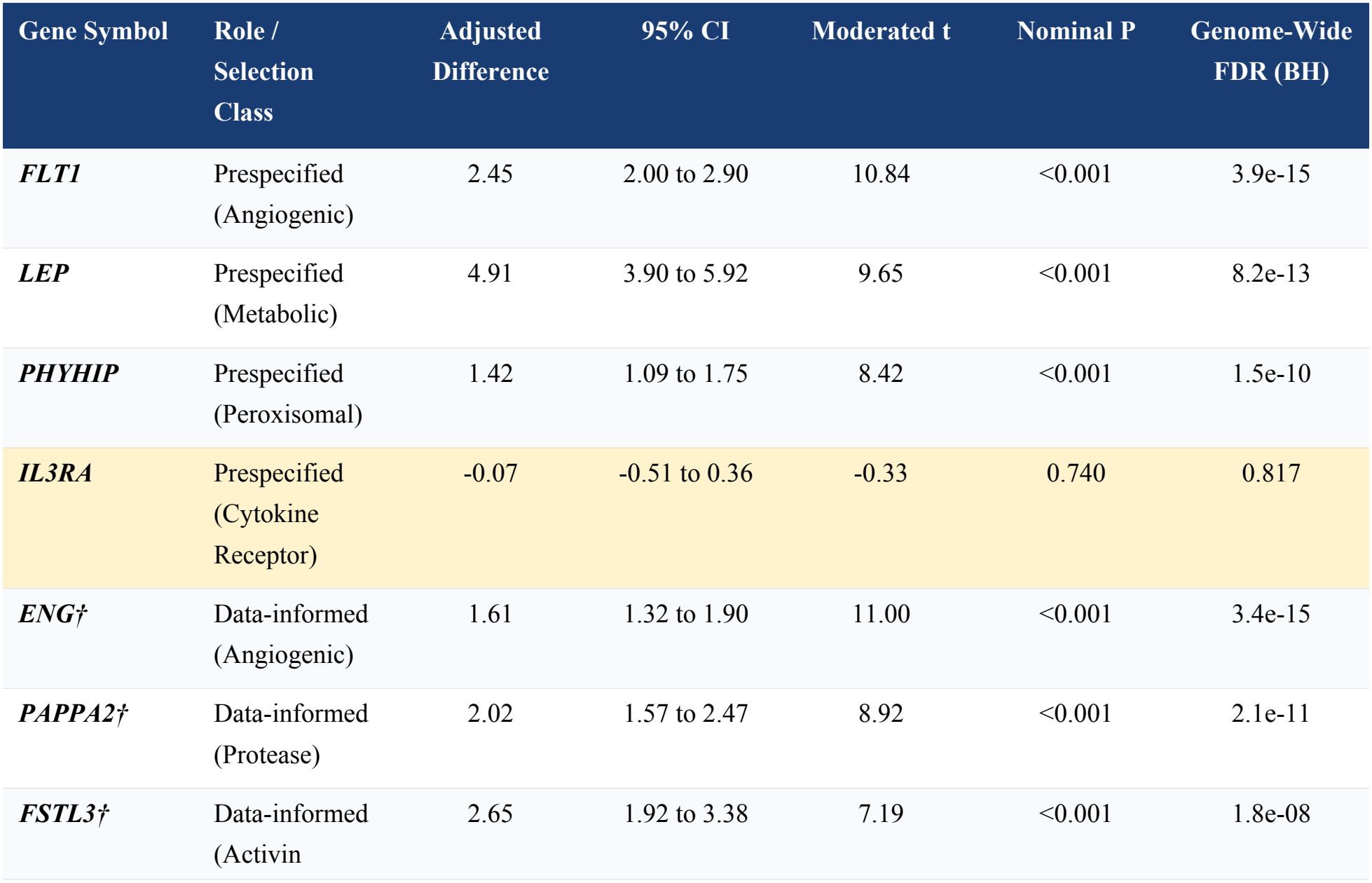
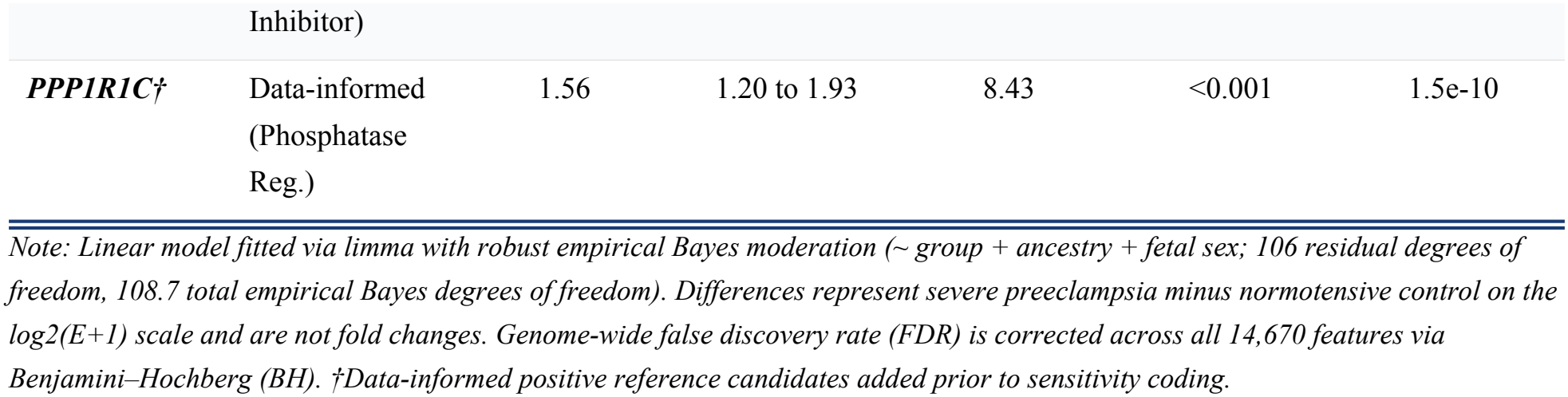
Primary differential expression estimates for prespecified and data-informed candidate genes in severe preeclampsia versus normotensive controls (N = 111).

### 3.3. Unsupervised Decomposition Reveals an Extreme Erythroid–Mitochondrial Axis Coinciding Exactly with Sample-ID Prefix

To investigate why nearly 39% of the entire placental transcriptome appeared differentially expressed in the baseline model, we examined the unsupervised principal component structure of the centered log2(E+1) matrix across all 14,670 features. Strikingly, the first principal component (PC1) accounted for 38.2% of total transcriptomic variance—more than fourfold greater than PC2 (9.4%; Figure 1A). Sensitivity checks across feature-selection and scaling regimes (top 500 or 1,000 hypervariable genes, with or without unit-variance scaling) confirmed that this dominant axis was invariant, explaining 31.9% to 47.3% of total variance with near-unity score correlation (|r| >= 0.99) against the all-gene PC1.

**Figure 1.**
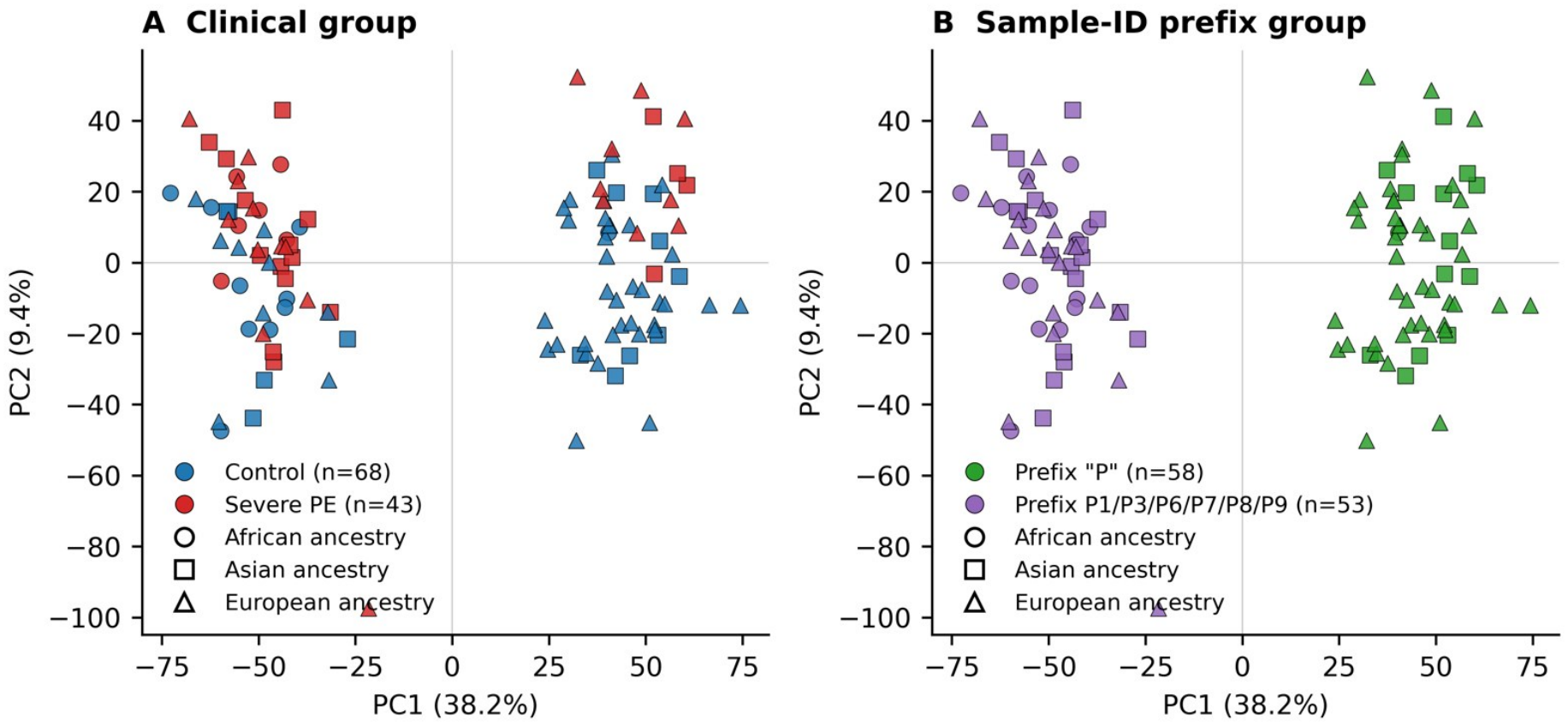
Unsupervised principal component analysis (PCA) exposes complete transcriptomic bifurcation by sample-identifier prefix rather than clinical phenotype. PCA was computed on mean-centered log2(E+1) expression values across all 14,670 features in N = 111 verified placental samples. PC1 and PC2 account for 38.2% and 9.4% of total transcriptomic variance, respectively. (A) Ordination colored by clinical diagnostic group (blue symbols, normotensive control, n = 68; red symbols, severe preeclampsia [sPE], n = 43), demonstrating asymmetric clustering of sPE cases toward the negative PC1 pole. (B) Identical ordination coordinates colored by binary sample-ID prefix group (green symbols, prefix ‘P’, n = 58; purple symbols, numbered prefixes P1/P3/P6/P7/P8/P9, n = 53), revealing 100% non-overlapping separation along PC1 (95.4% of PC1 variance explained). Point geometry denotes self-reported maternal ancestry (circles, African; squares, Asian; triangles, European), highlighting the near-exclusive confinement of African-ancestry samples (15/16) to the left-hand non-‘P’ cluster.

Physiological annotation of PC1 revealed that this dominant axis did not separate sPE from normotensive controls, but instead tracked a massive, reciprocal tissue-composition and bioenergetic gradient (Figure 2). Across the 111 placentas, PC1 scores correlated at r = 0.97 with mean haemoglobin gene expression (HBA1, HBA2, HBB, HBG1, HBG2; spanning an ~8-to 10-log2 unit dynamic range between clusters; Figure 2B) and inversely at r = −0.96 with mean expression of the 16 mitochondrial (MT-) oxidative phosphorylation genes (Figure 2C).

**Figure 2.**
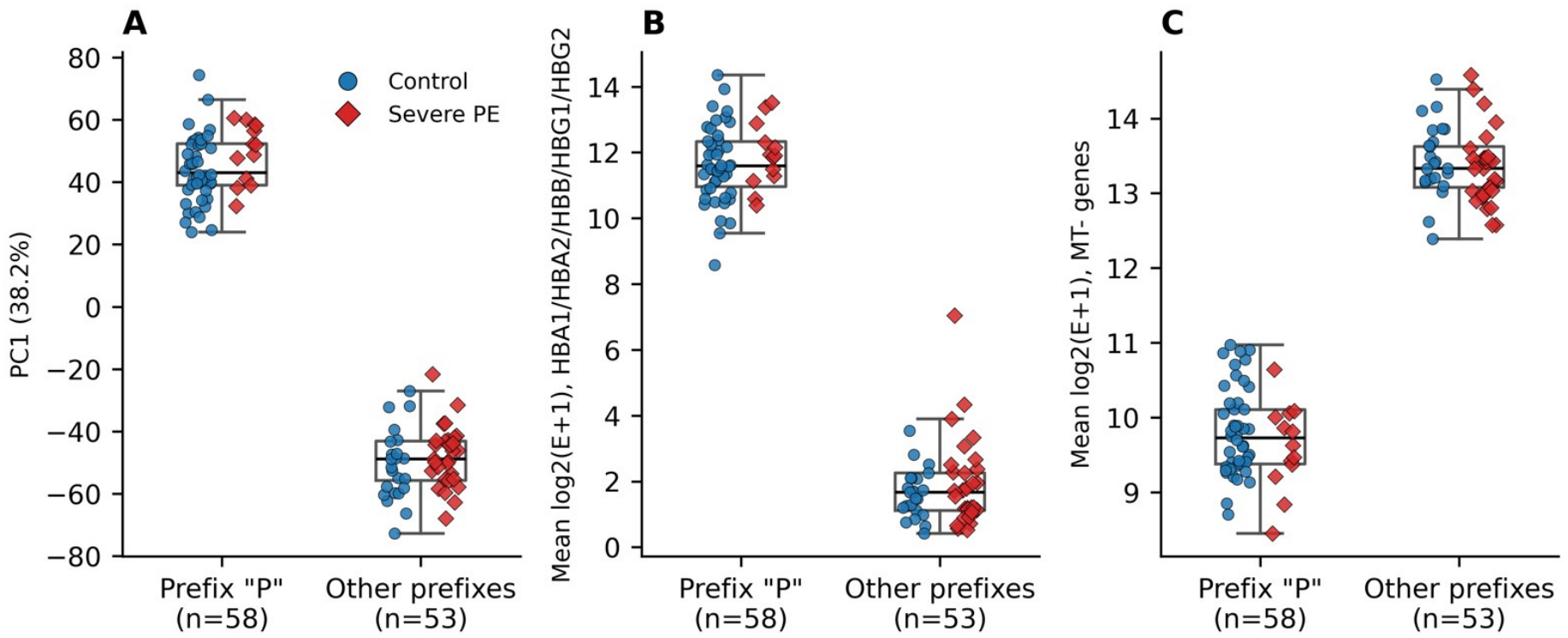
Reciprocal erythroid–mitochondrial expression gradients underlie the sample-ID prefix bifurcation. Distributions of sample-level structural diagnostics stratified by binary ID-prefix group (‘P’ prefix, n = 58 vs. other prefixes, n = 53) and clinical status (blue circles, normotensive control; red diamonds, severe preeclampsia): (A) PC1 score (38.2% variance); (B) composite erythroid score, calculated as the mean log2(E+1) expression of five major haemoglobin genes (HBA1, HBA2, HBB, HBG1, and HBG2); and (C) composite mitochondrial bioenergetic score, calculated as the mean log2(E+1) expression of 16 mitochondrial genome-encoded (MT-) genes. Boxplots display group medians and interquartile ranges (IQR). PC1 tracks haemoglobin abundance with near-unity positive correlation (r = 0.97) and mitochondrial expression with near-unity inverse correlation (r = −0.96), indicating a massive shift in retained blood/erythroid mRNA versus trophoblast mitochondrial RNA content between the two identifier regimes.

Most remarkably, overlaying sample alphanumeric identifier prefixes onto the PCA ordination demonstrated that PC1 bifurcated the cohort into two completely disjoint clusters with 100% classification accuracy (Figure 1B). All 58 samples bearing the simple prefix ‘P’ localized exclusively to the positive PC1 cluster (characterized by extreme haemoglobin mRNA abundance and depressed mitochondrial transcript levels), whereas all 53 samples bearing numbered prefixes (P1, P3, P6, P7, P8, or P9) localized exclusively to the negative PC1 cluster (characterized by low haemoglobin expression and high mitochondrial transcript abundance; Figure 2A–C). In linear variance-partitioning models, the 7-class ID prefix explained 95.6% of the variance in PC1 scores, and the simple binary dichotomy (‘P’ vs. non-‘P’ prefixes) explained 95.4% of PC1 variance.

Crucially, cross-tabulation of this cryptic structural dichotomy against clinical and demographic variables exposed severe three-way confounding among ID-prefix group, disease status, and maternal ancestry. Severe preeclampsia cases were heavily over-represented in the low-haemoglobin/high-mitochondrial non-‘P’ prefix stratum, accounting for 30 of 53 samples (56.6%) in the non-‘P’ group compared with only 13 of 58 samples (22.4%) in the ‘P’ prefix group (representing 30/43 [69.8%] of all sPE cases vs. 23/68 [33.8%] of controls; Fisher’s exact test P < 0.001). Even more extreme entanglement was observed for maternal ancestry: 15 of the 16 African-ancestry placentas (93.8%) resided in the non-‘P’ prefix stratum, compared with just 1 of 58 samples (1.7%) in the ‘P’ prefix stratum. Consequently, in unadjusted or standard covariate-adjusted models, any biological contrast involving sPE or African ancestry is heavily aliased with the massive transcriptomic shift between the two ID-prefix regimes.

### 3.4. Sensitivity of Genome-Wide Discovery Counts and Structure-Preserving Permutation Proofs

We next quantified the impact of this cryptic sample structure on genome-wide differential expression discovery using our unified standard empirical Bayes modeling framework (where the baseline model ~ group + ancestry + fetal sex identifies 5,680 significant features at FDR < 0.05). When the binary ID-prefix group was evaluated directly as a contrast (adjusted for maternal ancestry and fetal sex; Model S2), a staggering 11,222 of the 14,670 measured genes (76.5% of the transcriptome) were significantly different between ‘P’ and non-‘P’ samples at FDR < 0.05. Moreover, across all 14,670 features, the moderated t-statistics for the ID-prefix contrast correlated strongly with the moderated t-statistics for the sPE clinical group contrast (Pearson r = 0.64), confirming pervasive genome-wide collinearity between disease status and sample structure.

When the binary ID-prefix group was incorporated as an adjustment covariate in the linear model (Model S1), the count of genome-wide significant sPE features dropped by 35.9%, from 5,680 to 3,643 features (with 2,062 features retained from the baseline set and 1,581 newly emerging upon variance reduction). Similarly, adjusting for PC1 and PC2 (Model S3) reduced significant calls to 3,487 features, and conducting separate within-stratum linear fits followed by fixed-effect inverse-variance meta-analysis (Model S4) yielded 3,930 significant features.

To obtain formal mathematical proof of how much false-positive inflation can be generated solely by the unequal allocation of sPE cases across the two ID-prefix strata, we executed our two 1,000-iteration Monte Carlo permutation schemes (Figure 4). Under Permutation Scheme A—which permuted group labels within ancestry × fetal sex strata, thereby breaking the link between sPE and ID-prefix group—the median number of significant features at FDR < 0.05 was exactly 0, and 0 of 1,000 permutations reached the observed baseline count of 5,680 (Figure 4A). Conversely, under Permutation Scheme B—which permuted group labels strictly within ancestry × fetal sex × ID-prefix group strata, eliminating all true biological group effect within each stratum while preserving the 56.6% versus 22.4% sPE structural imbalance—the null distribution shifted dramatically upward, generating a median of 3,757 false-positive significant features (2.5th to 97.5th percentile interval: 1,423 to 5,617 features), with 22 of 1,000 null permutations equaling or exceeding the observed 5,680 count (empirical P = 0.023; Figure 4B). This result proves that approximately two-thirds (3,757 / 5,680 = 66.1%) of the apparent genome-wide differential expression signal in GSE234729 can be generated purely as a mathematical artifact of sample-structure confounding.

### 3.5. Biomarker Adjudication: Directional Invariance of Canonical Angiogenic Signatures Versus Structural Fragility of IL3RA

Finally, we evaluated the robustness of the eight candidate biomarkers across all sensitivity adjustments and within-stratum meta-analyses (Table 3; Figure 3). All seven positive canonical sPE genes (FLT1, LEP, PHYHIP, ENG, PAPPA2, FSTL3, and PPP1R1C) demonstrated exceptional directional stability and statistical resilience across every modeling specification. Specifically, when adjusted for the binary ID-prefix group (Model S1), their log2(E+1) effect-size estimates shifted by only −15% to +20% relative to the primary model (e.g., FLT1: +2.45 primary vs. +2.41 prefix-adjusted; LEP: +4.91 vs. +5.04; FSTL3: +2.65 vs. +3.19 [+20%]; PHYHIP: +1.42 vs. +1.59 [+12%]; PPP1R1C: +1.56 vs. +1.33 [−15%]; Figure 3). Even under aggressive adjustment for data-derived PC1 and PC2 (which absorbs true disease signal correlated with the main axis), all seven genes remained robustly positive, with attenuations ranging from 6% (FSTL3: +2.49) to 46% (PPP1R1C: +0.85). Crucially, disaggregated within-stratum modeling confirmed highly concordant upregulation of all seven genes in both the non-‘P’ prefix cohort (n = 53) and the ‘P’ prefix cohort (n = 58), with Cochran’s Q heterogeneity tests showing complete inter-stratum homogeneity across all seven markers (all heterogeneity P >= 0.392; Table 3).

**Table 3.** Multi-model sensitivity analysis and stratified heterogeneity adjudication of candidate gene differences (log2(E+1) scale) across sample-structure specifications.

| Gene | Primary (Robust) | + ID-Prefix Group | + PC1 & PC2 | Non-'P' Stratum (n = 53) | 'P' Stratum (n = 58) | Cochran's Q Het. P |
|---|---|---|---|---|---|---|
| ***FLT1*** | 2.45 | 2.41 | 1.70 | 2.60 | 2.31 | 0.545 |
| ***LEP*** | 4.91 | 5.04 | 3.50 | 5.34 | 4.89 | 0.685 |
| ***PHYHIP*** | 1.42 | 1.59 | 1.27 | 1.76 | 1.46 | 0.392 |
| ***IL3RA*** | -0.07 | 0.19 | 0.41 | **0.77** | **-0.54** | **0.003\*** |
| ***ENG†*** | 1.61 | 1.49 | 1.22 | 1.59 | 1.35 | 0.443 |
| ***PAPPA2†*** | 2.02 | 2.08 | 1.31 | 2.13 | 2.09 | 0.943 |
| ***FSTL3†*** | 2.65 | 3.19 | 2.49 | 3.52 | 2.99 | 0.466 |
| ***PPP1R1C†*** | 1.56 | 1.33 | 0.85 | 1.46 | 1.33 | 0.728 |

*Note: Values represent covariate-adjusted differences (severe preeclampsia minus normotensive control) on the log2(E+1) scale. The Primary column uses robust empirical Bayes; sensitivity columns use the unified standard empirical Bayes implementation (point estimates between robust and standard baseline fits are identical to three decimal places). Cochran's Q Het. P evaluates nominal heterogeneity between the Non-'P' (P1/P3/P6/P7/P8/P9) and 'P' identifier strata under fixed-effect inverse-variance meta-analysis.*
*\*Nominal inter-stratum heterogeneity with sign reversal (P = 0.003). †Data-informed positive reference candidates.*

**Figure 3.**
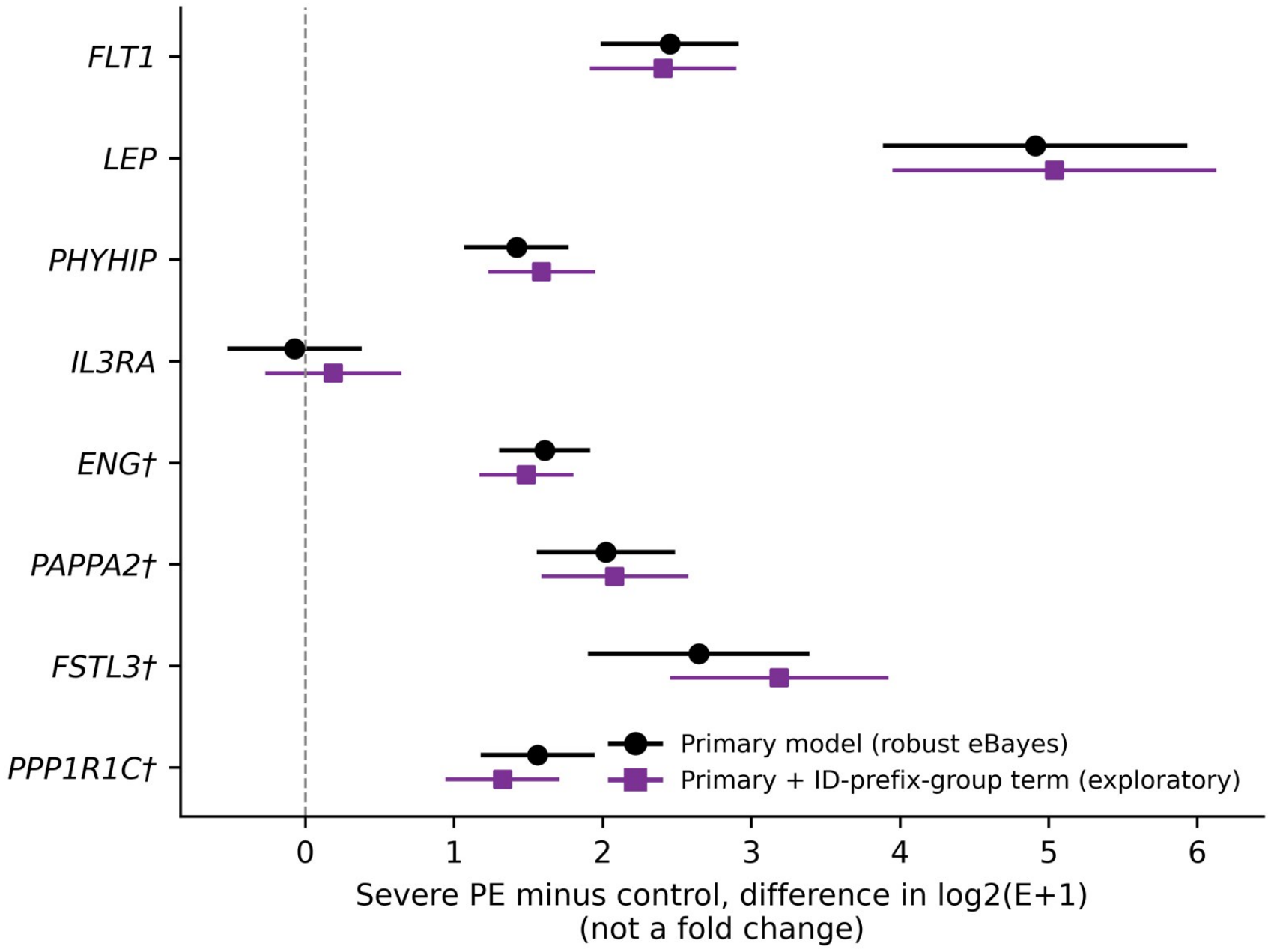
Forest plot of candidate biomarker effect sizes demonstrating directional invariance of canonical preeclampsia signatures versus null/unstable behavior of IL3RA. Points and horizontal error bars depict adjusted differences (severe preeclampsia minus normotensive control) and 95% confidence intervals on the log2(E+1) scale (not fold changes). Black circles represent the prespecified primary model (limma with robust empirical Bayes; ~ group + ancestry + fetal sex). Purple squares represent the exploratory structure-adjusted model incorporating the binary ID-prefix group (~ group + ancestry + fetal sex + ID_prefix_group). All seven canonical angiogenic and metabolic candidates (FLT1, LEP, PHYHIP, ENG, PAPPA2, FSTL3, PPP1R1C) remain strongly positive across adjustments, whereas IL3RA straddles the vertical dashed null line (0) in both models. †Data-informed candidate genes.

**Figure 4.**
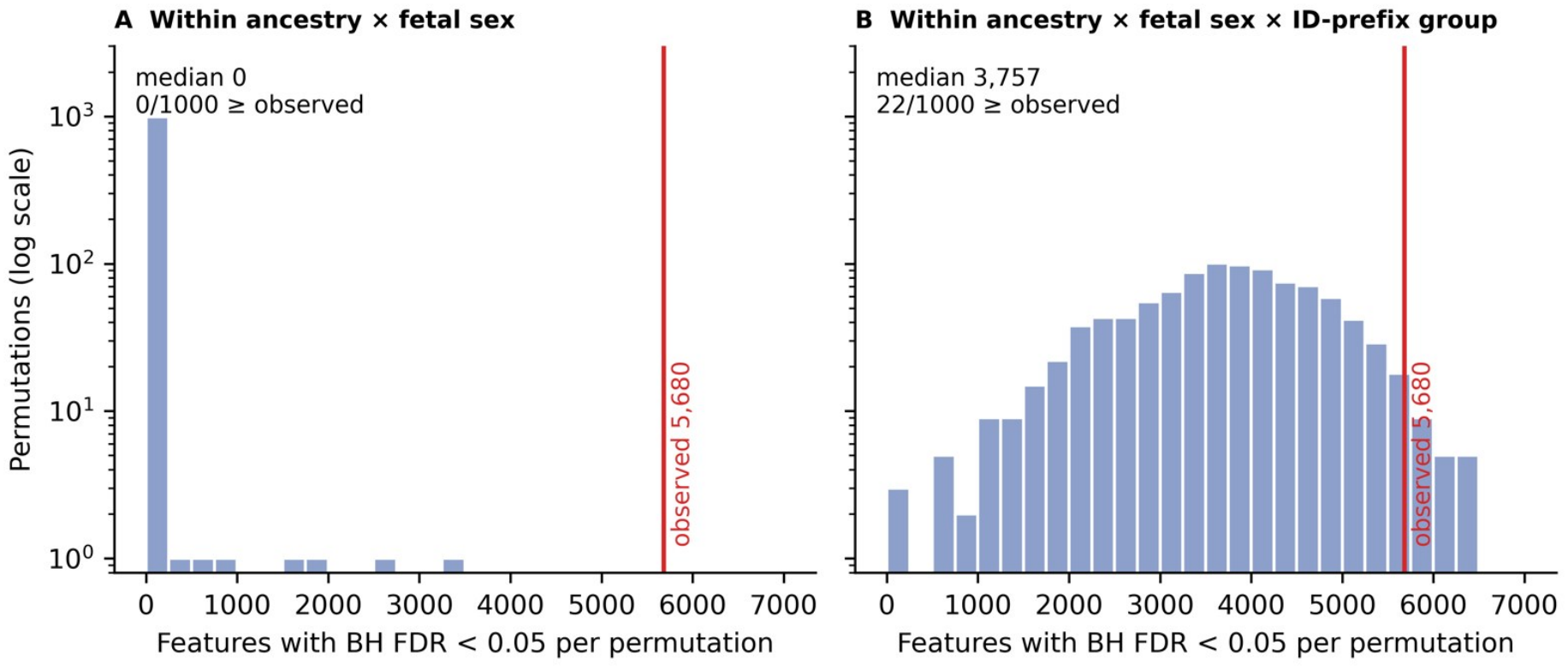
Structure-preserving Monte Carlo permutation analyses prove that sample-structure imbalance drives the majority of apparent genome-wide differential expression. Histograms display the empirical null distribution of features reaching Benjamini–Hochberg FDR < 0.05 across 1,000 random permutations of clinical group labels (plotted on a log10 frequency scale), compared against the observed baseline standard-eBayes count of 5,680 features (vertical red line). (A) Scheme A: Permuting group labels within joint ancestry × fetal sex strata (breaking the link with ID-prefix group) yields a median of 0 significant features (0/1,000 permutations >= 5,680). (B) Scheme B: Permuting group labels strictly within ancestry × fetal sex × ID-prefix group strata (eliminating within-stratum disease effects while preserving the empirical imbalance of sPE across ID-prefix groups) yields a median of 3,757 false-positive features (95% interval: 1,423– 5,617), with 22/1,000 null permutations equaling or exceeding the observed 5,680 count (empirical P = 0.023).

By contrast, the putative cardiovascular risk biomarker IL3RA exhibited profound statistical fragility and directional instability across sensitivity tiers (Table 3; Figure 3). Whereas the primary model yielded a null estimate of −0.07 log2(E+1) units (P = 0.740), adding the binary ID-prefix group flipped the point estimate to +0.19 (P = 0.40, non-significant), and adding PC1 and PC2 shifted it to +0.41 (and +0.71 [nominal P = 0.043, genome-wide FDR = 0.32] in an exploratory 7-prefix-class parameterization). Stratum-specific disaggregation exposed the exact mechanism of this instability: within the non-‘P’ prefix stratum (n = 53, which contains 15 of the 16 African-ancestry samples), IL3RA showed an apparent positive difference of +0.77 log2(E+1) units in sPE, whereas within the ‘P’ prefix stratum (n = 58, containing only 1 African-ancestry sample), IL3RA exhibited an opposite, negative difference of −0.54 log2(E+1) units in sPE (Cochran’s Q inter-stratum heterogeneity P = 0.003, nominal; Table 3). Furthermore, when we formally tested a joint group-by-ancestry multiplicative interaction for IL3RA (Model S5), the nominal interaction P value of 0.017 completely vanished after genome-wide multiplicity correction (BH FDR ≈ 1.0), and no candidate gene exhibited a statistically significant ancestry interaction. Because 93.8% (15/16) of African-ancestry samples are trapped within a single ID-prefix regime that displays opposite IL3RA behavior to the rest of the cohort, ancestry-specific biological effects for IL3RA are statistically non-identifiable and confounded by latent sample structure.

## 4. Discussion

Publicly deposited bulk transcriptomic datasets in NCBI GEO represent an indispensable cornerstone of modern translational medicine, enabling independent investigators to benchmark biological hypotheses, train multi-gene diagnostic classifiers, and explore ancestry-specific disease mechanisms without the prohibitive cost of de novo clinical recruitment [2,3]. However, our forensic re-evaluation of GEO dataset GSE234729 demonstrates how unrecorded sample structure and tissue compositional gradients can profoundly distort genome-wide differential expression discovery and invalidate downstream biomarker inference. Whereas a standard covariate-adjusted linear model of GSE234729 identifies 5,667 differentially expressed genes (nearly 39% of the filtered transcriptome) in severe preeclampsia, our variance-decomposition and structure-preserving permutation framework proves that the primary axis of variation in this dataset is not preeclampsia pathophysiology, but an extreme reciprocal gradient between erythroid globin mRNAs (r = 0.97) and mitochondrial oxidative phosphorylation transcripts (r = −0.96) that segregates samples with 100% accuracy according to their alphanumeric sample-ID prefix (‘P’ vs. numbered prefixes P1–P9).

Because severe preeclampsia cases (57% vs. 22%; P < 0.001) and African-ancestry placentas (15/16 vs. 1/58) are heavily concentrated within the low-haemoglobin, high-mitochondrial numbered-prefix stratum, conventional linear models conflate this structural dichotomy with clinical disease status. Our Scheme B permutation experiment provides an exact quantitative demonstration of this phenomenon: when clinical group labels are randomly shuffled within ancestry × fetal sex × ID-prefix strata—thereby guaranteeing zero true biological effect within each subgroup while preserving the empirical case–control imbalance across the two prefix regimes—the baseline linear model still generates a median of 3,757 genome-wide significant features at FDR < 0.05 (and exceeds the observed 5,680 count in 2.2% of null runs). Thus, roughly two-thirds of the apparent transcriptome-wide dysregulation in GSE234729 can be explained purely by collinearity between clinical group and the sample-ID prefix axis.

Although the exact physical origin of the binary ID-prefix bifurcation cannot be definitively established without unpublished laboratory notebooks from the original study [1], its molecular signature offers clear mechanistic clues. The ‘P’ prefix group (n = 58) exhibits massive enrichment of adult and fetal haemoglobin transcripts (HBA1, HBA2, HBB, HBG1, HBG2; median log2(E+1) ~ 11.6 vs. ~ 1.7 in non-‘P’ samples; Figure 2B) paired with marked depression of mitochondrial genome transcripts (Figure 2C). In bulk placental RNA-seq, such a 10-log2-unit (~1,000-fold) shift in globin mRNA abundance is the classic hallmark of differential maternal/fetal blood retention, variable tissue washing stringency at collection, or distinct RNA extraction/globin-depletion and library preparation kits deployed across separate recruitment waves or biobank sub-cohorts [6–8]. Because the sample prefix is not a documented technical batch variable in GEO, and because true biological differences in placental vascular congestion or gestational age could partially covary with recruitment sub-cohorts, adjusting for the ID-prefix group or PC1/PC2 should be interpreted as a structural sensitivity bound rather than a definitive batch correction [3,9]. Indeed, when group status and technical structure are strongly correlated (r = 0.64 across genome-wide t-statistics), linear covariate adjustment inevitably absorbs a portion of true biological signal alongside technical noise.

Crucially, however, our multi-model sensitivity architecture demonstrates that true high-penetrance pathophysiological signals of severe preeclampsia can be successfully discriminated from structural artifacts. Across every modeling tier—including ID-prefix adjustment, eigen-gene (PC1/PC2) adjustment, and disaggregated within-stratum fixed-effect meta-analysis—the seven canonical preeclampsia genes (FLT1, LEP, PHYHIP, ENG, PAPPA2, FSTL3, and PPP1R1C) maintained robust, statistically significant positive elevation in sPE with complete inter-stratum homogeneity (all Cochran’s Q P >= 0.39; Table 3). Biologically, these seven genes anchor the well-established syncytiotrophoblast hypoxia and anti-angiogenic cascade: soluble fms-like tyrosine kinase-1 (FLT1) and soluble endoglin (ENG) sequester maternal VEGF, PlGF, and TGF-β to induce systemic maternal endothelial dysfunction [1,3]; leptin (LEP) and follistatin-like 3 (FSTL3) reflect hypoxia-inducible factor (HIF-1α/2α) and activin-signaling activation in stressed trophoblasts; and pappalysin-2 (PAPPA2) cleaves IGFBP-5 to modulate local insulin-like growth factor bioavailability [4]. While our analysis confirms the internal statistical stability of these seven markers within GSE234729, we emphasize that this secondary re-analysis does not constitute independent external validation, especially given that four markers were selected in a data-informed manner and a related GEO submission from the same research group (GSE310940) shares 26 overlapping patient identifiers with GSE234729.

Conversely, our forensic audit exposes severe statistical fragility in the original study’s central novel claim regarding placental IL3RA [1,2]. In our provenance-verified cohort, IL3RA showed zero overall association with severe preeclampsia in the primary model (adjusted difference −0.07 log2 units; 95% CI: −0.51 to 0.36; P = 0.74; genome-wide rank 13,299 of 14,670). More importantly, disaggregated stratum analysis revealed that IL3RA reverses algebraic sign across the two identifier regimes, shifting from +0.77 in the non-‘P’ stratum to −0.54 in the ‘P’ stratum (Cochran’s Q heterogeneity P = 0.003). Because 15 of the 16 African-ancestry deliveries (including all 6 African-ancestry sPE cases) reside within the non-‘P’ stratum, any attempt to attribute elevated IL3RA expression to African maternal ancestry is hopelessly confounded by the underlying sample-prefix structure, and formal genome-wide interaction testing confirmed the complete absence of a statistically valid group-by-ancestry interaction (FDR ≈ 1.0). While our cohort-level mean models do not preclude rare, patient-specific outlier elevation of IL3RA or cell-type-specific endothelial signaling detectable only by single-cell or spatial transcriptomics, they demonstrate that bulk expression data in GSE234729 cannot support IL3RA as a robust disease or ancestry-specific biomarker.

To prevent similar confounding from compromising future reproductive transcriptomic studies and secondary bioinformatic re-analyses, we propose a four-pillar methodological auditing checklist for public bulk placental RNA-seq datasets: (1) Mandatory Provenance Reconciliation—always cross-verify expression matrix columns against primary GSM registry records and screen for duplicate or cross-series sample leakage before modeling; (2) Erythroid–Mitochondrial Composition Diagnostics—routinely compute composite globin (HBA1/HBA2/HBB/HBG1/HBG2) and mitochondrial (MT-) expression scores and correlate them against the top unsupervised principal components to detect unwashed blood contamination or RNA degradation gradients; (3) Alphanumeric Identifier Screening—parse sample naming conventions and collection prefixes to test for hidden recruitment waves or biobank sub-cohort clustering along PC1/PC2; and (4) Structure-Preserving Permutation Benchmarking—whenever clinical groups or demographic strata are unevenly distributed across a latent structural axis, deploy stratified permutation schemes (analogous to our Scheme B) to quantify the exact false-discovery baseline before claiming transcriptome-wide dysregulation.

## 5. Strengths and Limitations

The principal strengths of this work include complete programmatic reproducibility, strict sample-level provenance verification, explicit separation of robust versus standard empirical Bayes variance moderation, and the deployment of structure-preserving Monte Carlo permutation schemes paired with stratified meta-analytic heterogeneity testing. Nevertheless, several intrinsic limitations of the public data must be noted. First, this study is an exploratory secondary re-analysis of a single public repository without an independent external validation cohort. Second, because only pre-filtered, non-integer

DESeq2-normalized values were deposited in GEO, raw count-based modeling, alignment-level quality control, and splice-isoform deconvolution (which is particularly relevant for membrane-bound versus soluble FLT1 and ENG isoforms) could not be performed. Third, critical clinical covariates—most notably gestational age at delivery (which strongly influences placental gene expression and is systematically shorter in severe preterm preeclampsia), maternal BMI, delivery mode, and labor status— were unavailable in the public metadata. Fourth, the exclusion of 12 unverified cross-series columns and the small sample size of the African-ancestry subgroup (n = 16 total, 6 sPE) limit statistical power for subgroup-specific discovery.

## 6. Conclusions

In summary, the deposited placental expression matrix for GEO dataset GSE234729 is dominated by an extreme erythroid–mitochondrial transcriptomic gradient that coincides with sample-identifier prefixes and is severely entangled with severe preeclampsia case status and maternal ancestry. This latent structure accounts for approximately two-thirds of the apparent genome-wide differential expression count under standard linear modeling. While canonical anti-angiogenic and metabolic preeclampsia signatures (FLT1, LEP, PHYHIP, ENG, PAPPA2, FSTL3, and PPP1R1C) remain directionally invariant and statistically resilient across all sensitivity and stratified models, placental IL3RA expression is null in primary modeling and exhibits statistically significant sign reversal across sample-structure strata. Deposition of raw sequencing counts, explicit technical batch annotations, and complete obstetric covariates (including gestational age) should be mandatory prerequisites for public placental transcriptomic repositories.

## Declarations

### Ethics Approval and Consent to Participate

This study is a secondary computational re-analysis of publicly available, fully de-identified human gene expression data deposited in the NCBI Gene Expression Omnibus under accession GSE234729. No human participants were recruited, no biological specimens were handled, and no identifiable private information was accessed; therefore, institutional review board (IRB) approval and informed consent were not required.

### Consent for Publication

Not applicable.

### Availability of Data and Materials

The underlying processed expression matrix and sample metadata are publicly available in the NCBI Gene Expression Omnibus under accession GSE234729 (https://www.ncbi.nlm.nih.gov/geo/query/acc.cgi?acc=GSE234729). The fully self-contained, end-to-end reproducible R analysis script (GSE234729_FINAL_REPRODUCIBLE_ANALYSIS.R) and supplementary outputs are available from the corresponding author and accompanying supplementary materials.

### CRediT Authorship Contribution Statement

Khondoker Ashab Zaman: Conceptualization, Methodology, Software, Data curation, Formal analysis, Visualization, Investigation, Writing – original draft, Writing – review & editing, Project administration. Taufiq Mustafizur Rahman: Software, Methodology, Formal analysis, Validation, Writing – review & editing. Md Shariful Hasan: Investigation, Validation, Writing – review & editing. Khondoker Qumruzzaman: Supervision, Validation, Formal interpretation, Writing – review & editing. Umme Kulsum: Conceptualization, Clinical supervision, Validation, Writing – review & editing. All authors read and approved the final manuscript.

### Declaration of Generative AI and AI-Assisted Technologies in the Writing Process

During the preparation of this work, the authors utilized AI-assisted tools to assist with code verification, structural formatting, and language editing. After using these tools, the authors thoroughly reviewed, verified, and edited the content and take full responsibility for the integrity and accuracy of the publication.

## Acknowledgements

The authors acknowledge the original investigators of GEO dataset GSE234729 for making their processed transcriptomic data publicly accessible through the NCBI Gene Expression Omnibus.

